# AP2A1 is upregulated upon replicative senescence of human fibroblasts to strengthen focal adhesions via integrin β1 translocation along stress fibers

**DOI:** 10.1101/2023.08.19.553998

**Authors:** Pirawan Chantachotikul, Shiyou Liu, Kana Furukawa, Shinji Deguchi

## Abstract

Aging proceeds with accumulation of senescent cells in multiple organs. Senescent cells become large in size compared to young cells, which promotes further senescence and age-related diseases. Currently, the molecular mechanism behind the maintenance of such huge cell architecture undergoing senescence remains poorly understood. Here we focus on reorganization of actin stress fibers induced upon replicative senescence of human fibroblasts, typically used as a senescent cell model. We identified, together with our previous proteomic study, that AP2A1 (alpha 1 adaptin subunit of the adaptor protein 2) is upregulated in senescent cells along the length of stress fibers, which are enlarged following the increase in the whole cell size. We then revealed that knockdown of AP2A1 in senescent cells suppresses key senescence-associated phenotypes, which include decreased cell area and lowered expression of major senescence markers. Meanwhile, AP2A1 overexpression in young cells induced the opposite effects that rather advance senescence, suggesting that AP2A1 may be used as a senescence marker. We found that AP2A1 is colocalized with integrin β1, and both of them move linearly along stress fibers. We further observed that focal adhesions are enlarged in senescent cells to reinforce cell adhesions to the substrate. These results suggest that senescent cells maintain their large size by strengthening the anchorage to the substrate by supplying integrin β1 via translocation along stress fibers. This mechanism may work efficiently in senescent cells, compared with a case relying on random diffusion of integrin β1, given the enlarged cell size and resulting increase in travel time and distance for endocytosed vesicle transportation.

## 1. Introduction

Accumulation of senescent cells in multiple organs during aging contributes to age-associated diseases (Phillip et al., 2015; Valentijn et al., 2018), which include neurodegenerative diseases (Kritsilis et al., 2018; Martinez-Cué & Rueda, 2020), cardiovascular diseases (Childs et al., 2018; Olivieri et al., 2013), type 2 diabetes (Palmer et al., 2019; Shakeri et al., 2018), and cancer (Campisi, 2013; Krtolica et al., 2001). Cultured human fibroblasts have been used in the study of cellular senescence given that they display a limited capacity for cell division before entering a stable proliferative growth arrest, a state known as “replicative senescence” (Hayflick & Moorhead, 1961). The replicative senescence is caused by a telomere shortening upon each cell division, resulting in the accumulation of DNA damage (Courtois-Cox et al., 2008). Advanced senescence gradually leads to an increase in cell size (Narita et al., 2003), which in turn impairs cell function, proliferative ability, and protein synthesis (Neurohr et al., 2019). Senescence is also accompanied by elevated activity of senescence-associated β-galactosidase (SA-β-gal) and expression of cell cycle inhibitors such as p53, p21, p16, and PAI-1 (Calcinotto & Alimonti, 2017). While cell morphological change is thus commonly recognized as a senescence indicator, the underlying mechanisms remain largely unknown. Replicative senescence also causes a decrease in cell migratory potential (Geissler et al., 2012; Younis et al., 2018). In this regard, the accumulation of senescent cells leads to poorly healing wounds (Mulder & Vande Berg, 2002; Stanley & Osler, 2001; Telgenhoff & Shroot, 2005).

The regulation of cell morphology and migration is closely associated with the dynamics of stress fibers (Kassianidou & Kumar, 2015; Vicente-Manzanares et al., 2008). Stress fibers are actomyosin-based bundles, which are composed mainly of actin filaments cross-linked by α-actinin and non-muscle myosin II (Burridge & Wittchen, 2013; Cramer et al., 1997; Lazarides & Burridge, 1975). In mesenchymal cell types, stress fibers localize between the adhesion sites to generate contractile force on the underlying extracellular matrix (Burridge & Guilluy, 2016; Li et al., 2022). Changes in stress fiber organization and contractile properties affect numerous cellular processes including adhesion (Geiger et al., 2009; Parsons et al., 2010), motility (Chen, 1981; Vicente-Manzanares et al., 2009), and mechanosensing (Burridge & Wittchen, 2013; Huang et al., 2021). The organization of stress fibers, known to be altered in cells undergoing senescence (Chen et al., 2000; Moujaber et al., 2019), has also been reported to play a role in pathological processes including developmental morphogenesis and cancer metastasis (Ridley et al., 2003; Tojkander et al., 2012).

While the proteome of stress fibers has only been partly investigated, our group recently revealed that those in human fibroblasts are comprised of at least 135 proteins, and 63 of them are upregulated with replicative senescence (Liu et al., 2022). Approximately one-third of the upregulated proteins belongs to the actin cytoskeletal component that includes caldesmon, tropomyosin, and filamin, which are all well characterized and are indeed likely to be enriched in the aged thick stress fibers; meanwhile, the function of the rest related to stress fibers remains unclear. Among them, here we focus on AP2A1 (alpha 1 adaptin subunit of the adaptor protein 2) because its molecular mechanisms behind the link to stress fiber regulation and cellular senescence have not been reported. AP2A1 is known to be a major protein involved in vesicle formation in clathrin-mediated endocytosis (CME) by recognizing and binding cargo proteins, clathrin, and accessory proteins (Di Rubbo et al., 2013; Kadlecova et al., 2017; Shin et al., 2021). AP2A1 has also been implicated in many diseases such as skin and neurodegenerative disorders (Wang et al., 2017), but molecular details remain unclear. Using human fibroblasts undergoing replicative senescence, here we found that the expression level of AP2A1 controls the extent of the senescence progression, specifically, e.g., the expression of senescence markers, morphological and migratory phenotypes, and thickness and turnover of individual stress fibers. We then showed that AP2A1 plays a role in integrin β1 translocation along the length of stress fibers, which is enhanced in aged cells to strengthen cell adhesions. Taken together, our study uncovered the mechanisms behind the morphological alteration of stress fibers upon senescence and proposes AP2A1 as a useful biomarker for assessing cellular senescence.

## 2. Materials and Methods

### 2.1 Cell culture

Primary human foreskin fibroblasts (HFF-1, ATCC) were cultured as a monolayer in high-glucose Dulbecco’s Modified Eagle’s Medium (DMEM, Wako) supplemented with 15% fetal bovine serum (Sigma-Aldrich) and 1% penicillin-streptomycin solution (Wako). Cells were kept at 37°C in a humidified incubator with 5% CO_2_. The medium was changed once every 3 days, and cells were passaged at a split ratio of 1:3 when 80% confluence was reached.

### 2.2 SA-β-gal staining

Cell senescence was assessed by the senescence-associated β-galactosidase staining kit (Cell Signaling Technology) following the manufacturer’s protocol. Briefly, cells at passage 10, 20, and 30 (hereafter described as P10, P20, and P30, respectively) were grown in 6-well plates until sub-confluence. Cells were fixed in 4% paraformaldehyde in phosphate-buffered saline (PBS, Wako) at 37°C after the media were removed and then were incubated with the β-galactosidase staining solution (pH 6.0) at 37°C overnight in a dry incubator. Images were taken at 10 random fields using an inverted microscope (Olympus CKX41) to determine the percentage of SA-β-gal positive cells stained in blue.

### 2.3 Cell proliferation assay

Cell proliferation was assessed using the 5-Ethynyl-2-deoxyuridine (EdU) incorporation assay (Click-iT EdU Alexa Fluor 488 Imaging Kit, Invitrogen) following the manufacturer’s protocol. Briefly, cells were seeded on glass bottom dishes (Ø 27 mm, No. 1S, Iwaki) and treated with 20 *µ*M EdU solutions diluted in culture media for 24 h at 37°C. Cells were then fixed with 4% paraformaldehyde, permeabilized with 0.5% Triton X-100 in PBS for 20 min, and treated with Cu(I)-catalyzed click reaction cocktails for 30 min. Cells were also stained with Hoechst 33342 (1∶2000) in PBS. Stained cells were observed under a confocal laser scanning microscope (FV1000, Olympus) equipped with a UPlan Apo 60x oil objective lens (NA = 1.42). EdU-stained cells (proliferative cells) and Hoechst 33342-stained cells (total cells) were counted to evaluate the ratio of proliferative cells to the total cells.

### 2.4 Quantification of protein synthesis

To quantify protein synthesis, the Click-iT^TM^ HPG Alexa Fluor 488 Protein Synthesis Assay Kit was used following the manufacturer’s protocol, which evaluates the incorporation of L-homopropargylglycine (HPG) to newly synthetized proteins. Briefly, cells were incubated in methionine-free DMEM media for 1 h to deplete endogenous methionine and then in other methionine-free DMEM media containing 2 mM HPG for 30 min to allow incorporation of HPG to nascent proteins. Cells were then fixed and permeabilized for fluorescence imaging. The incorporation of HPG, quantified by the intensity of Alexa Fluor 488, was defined in each cell as the mean intensity of the pixels inside the cell boundaries after background subtraction.

### 2.5 Cellular ATP measurement

Intracellular ATP levels were measured using the ATP assay Kit-Luminescence (Dojindo Molecular Technologies) following the manufacturer’s protocol. Briefly, 5,000 cells in a well of 96-well plates were lysed with the lysis buffer of 50 *µ*l and incubated at 25°C for 10 min. Luminescence signals from the ATP standard solution and samples were detected on a microplate reader (Spectra MAX GeminiEM, Molecular Devices).

### 2.6 Immunostaining

Cells were seeded at a low density on coverslips, fixed with 4% paraformaldehyde in PBS for 15 min, permeabilized with 0.5% (V/V) Triton X-100 for 15 min, and blocked with 1% (w/v) normal goat serum in PBS for 1 h at room temperature. Cells were then incubated overnight at 4°C with primary antibodies, anti-AP2A1 (1:200, #MA1-064; Thermo Fisher Scientific), anti-vinculin (1:200, #ab11194; Abcam), anti-paxillin (1:200, #610569, BD Biosciences), anti-integrin β1 (1:200, #26918-1-AP; Proteintech Group), and anti-lysosomal-associated membrane protein-1 (LAMP-1, 1:200 #14-1079-80; Thermo Fisher Scientific), followed by incubation with secondary antibodies (anti-rabbit conjugated with Alexa Flour 488, anti-rabbit conjugated with Alexa Flour 546, anti-mouse conjugated with Alexa Flour 546; Invitrogen) for 2 h at 37°C. To visualize stress fibers, cells were stained with AlexaFluor™ 488 phalloidin (ActinGreen 488 ReadyProbes Reagent; Thermo Fisher Scientific) or phalloidin 647 (Funakoshi) for 20 min at room temperature. Finally, cell nuclei were stained with Hoechst33342 (1 *µ*g/ml final concentration), and samples were mounted with ProLong Gold Antifade Mountant (P36930, Thermo Fisher Scientific). Laser scanning confocal microscope (Olympus FV3000) equipped with a UPlan Apo 60x oil objective lens (NA = 1.42) or structured illumination microscopy (SIM, Zeiss Elyra S.1; Center for Medical Innovation and Translational Research (COMIT) of Osaka University Medical School) equipped with an alpha Plan-Aprochromat 100x oil objective lens (NA = 1.46) were used for imaging.

### 2.7 Quantification of cell area

The captured fluorescence images were imported to Fiji/ImageJ software. After setting the scale, the cell contour was outlined by function “Polygon Selection”, and then cell areas were measured by function Measure. For analyzing the average of the cell areas, data measured from a total of 90 cells captured in each experimental condition were averaged.

### 2.8 Quantification of stress fiber thickness

Grey-scale images were used for fiber segmentation, which was performed in Fiji/ImageJ software by subtraction of non-uniform background within the cell boundaries. Line region of interest (ROI) was used to draw a ROI across the width of individual stress fibers. Two-dimensional graphs showing the intensities along the line selection were displayed to obtain the horizontal distance between the two edges of the stress fibers. The thickness of stress fibers was then determined as [(Peak intensity) – (Valley intensity)]/2. A total of 300 measurements were performed.

### 2.9 Fiber alignment analysis

Stress fibers and AP2A1 were imaged with confocal microscopy (Olympus FV3000) to evaluate their alignment using two-dimensional fast Fourier transform (FFT) in Fiji/ImageJ software. The FFT images provide a frequency map, in which the direction of the structure alignment is detected. Image components of high and low frequency, corresponding to short and long periodicity, are represented as peaks around the periphery and center in the Fourier spectrum, respectively. A circular projection was drawn at the center of the FFT images, and Oval Profile plugin was used to determine the direction based on the pixel intensity along the radial direction. The degree of alignment was represented by the height and shape of the pixel intensity plot.

### 2.10 Colocalization analysis

To analyze colocalization of endogenous proteins, specifically actin filaments, AP2A1, and transferrin, raw images (resolution: 1024 × 1024 pixels) captured by confocal microscopy (Olympus FV3000) were processed in JACoP plugin of Fiji/ImageJ software. Cells were identified in manually selected ROI. Pearson correlation coefficient was calculated to determine the correlation between two fluorescence signals.

### 2.11 Wound healing analysis

Cells were seeded in 12-well plates and incubated for 24 h until they reached 90% confluence. A 200-μl sterile pipette tip was used to scratch the cells, and detached cells were removed with PBS. The plate was placed on a phase-contrast microscope (Olympus IX-73) to capture the resulting cell migration using a 10x objective every 30 min for 12 h. We determined the percentage of wound closure, namely the rate of cell migration, in Fiji/ImageJ software.

### 2.12 Transferrin uptake assay

Cells were grown on coverslips, pre-incubated with Opti-MEM (Gibco) for 2 h, incubated with Alexa-Fluor-488-conjugated transferrin of 25 *µ*g/ml (Thermo Fisher Scientific) in Opti-MEM for 10 min on ice, shifted to 37°C for 15 min to allow uptake of transferrin, washed with PBS twice, and fixed with 4% formaldehyde. The fluorescence images of cells were obtained using a confocal microscopy (Olympus FV3000) and analyzed with the Fiji/ImageJ software to evaluate the transferrin uptake in individual cells, which were selected as ROI by using a polygon tool.

### 2.13 Isolation of stress fibers

Stress fibers were isolated from cells as previous reported (Deguchi et al., 2012; Liu et al., 2022; Matsui et al., 2011; Okamoto et al., 2020) Briefly, cells grown on 100-mm polystyrene cell culture dishes were washed twice with ice-cold PBS, hypotonically shocked with a low-ionic-strength extraction solution (2.5mM triethanolamine, 1 mM dithiothreitol, 1μg/ml pepstatin, and 1 μg/ml leupeptin in ultra-pure water) for 5 min, incubated with 0.05% Triton X-100 in the cytoskeleton stabilizing buffer (20 mM Imidazole, 2.2 mM MgCl2, 2 mM EGTA, 13.3 mM KCl, 1 mM dithiothreitol, 1 μg/ml pepstatin, and 1 μg/ml leupeptin; pH 7.3) for 1 min, and rinsed with the same cytoskeleton stabilizing buffer without Triton X-100. The extracted stress fibers were scraped off from the dish, suspended in lysis buffer (50 mM Tris-HCl, 100 mM NaCl, 1% NP-40, 1% sodium deoxycholate, 1 mM Na_3_VO_4_, 1 mM NaF, 1 mM dithiothreitol, 1 mM phenylmethylsulfonyl fluoride, 1μg/ml pepstatin, and 1 μg/ml leupeptin; pH 7.4). The stress fiber lysates were cleared by centrifugation (15,000 x g, 60 min) and were kept on ice for western blot analysis.

### 2.14 Western blotting

To prepare the whole cell extracts, cells were washed twice with ice-cold PBS and scraped off using a plastic scraper from the culture dish containing lysis buffer. The lysates from the extracted stress fibers or whole cells were maintained on a shaker for 60 min at 4°C and centrifuged at 15,000 x g for 60 min to collect the supernatant. The total protein concentration was determined by Pierce BCA Protein Assay Kit (Thermo Fischer Scientific). Proteins were fractionated by 10% SDS-PAGE and transferred onto polyvinylidene fluoride membranes (0.45 μm, Wako). The membranes were blocked in 5% (w/v) BSA for 1 h at room temperature, followed by incubation overnight at 4°C with primary antibodies: anti-GAPDH (1:5000; #014-25524; Wako), anti-p53 (1:1000; #10442-1-AP; Proteintech), anti-p21 (1:1000; #10355-1-AP; Proteintech), anti-myosin IIa (1:1000; #PRB-440P; Covance), anti-α-actinin (1:1000; #ab18061; Abcam), anti-β-actin (1:1000; #4970P; Cell Signaling Technology), and anti-AP2A1 (1:1000; #11401-1-AP; Proteintech). Excess antibodies were removed by washing with TBS-T three times. Membranes were then incubated with HRP-conjugated anti-rabbit/mouse for 1 h. After 3 washes with TBS-T, immunoreactive proteins were detected using Immobilon Western Kit (Millipore). Protein bands from each blot were observed by an imaging system (ChemiDoc XRS+, Bio-Rad) and analyzed by ImageLab software (Bio-rad). The GAPDH was used as an internal control.

### 2.15 Plasmids, transfection, and knockdown

An expression vector encoding mClover2-human AP2A1 was constructed from the pEX-A2-AP2A1 vector (Eurofins Genomics), which was digested with EcoRI and BamHI restriction enzymes and cloned into the EcoRI/BamHI-double-digested mClover2-C1 vector (Addgene plasmid#54577). Plasmid vectors for expressing human actin binding protein Lifeact (mRuby2-tagged Lifeact; Saito et al., 2021) and EGFP-tagged myosin regulatory light chain (MLC; Huang et al., 2021) were generated as previously described. mVenus-Integrin-Beta1-N-18 plasmid (Addgene plasmid #56330) was obtained from Addgene (USA). Transfection was performed using Lipofectamine LTX and Plus reagent (Invitrogen Life Technologies) following the manufacturer’s instructions with plasmids of 2.5 *µ*g per well of 35-mm culture dishes where cells were grown to 50–70% confluence in Opti-MEM medium. For co-transfection, the same amounts of two types of plasmids were applied to make 2.5 *µ*g in total. Cell imaging was performed 24 h after transfection. The siRNAs targeting AP2A1 (s183 and s184, Thermo Fisher Scientific) and negative control siRNAs (Cell Signaling Technology) were transfected to cells to evaluate the effect of downregulating the expression of AP2A1. siRNA transfection was carried out at a concentration of 60 pmol in Opti-MEM medium using Lipofectamine RNAi-MAX (Invitrogen Life Technologies) following the manufacturer’s instructions, and then cell imaging was performed 48 h after the transfection.

### 2.16 Fluorescence recovery after photobleaching (FRAP)

Fluorescence recovery after photobleaching (FRAP) was performed by using the laser scanning microscope (Olympus FV3000). Images were acquired using a 60x oil immersion objective (NA = 1.42) with line scanning. Two images were acquired to monitor the fluorescence signal before bleaching, and square areas of EGFP-MLC-labeled stress fibers were then photobleached using 405 and 488-nm wavelength lasers. The subsequent fluorescence recovery was recorded at a 20-sec interval for 6 min.

### 2.17 Time-lapse imaging of protein transport

Cells at P10 and P30 were cotransfected with the two constructs: mClover2-AP2A1 (green) and mRuby2-tagged Lifeact (red) or mVenus-Integrin-Beta1 (green) and mRuby2-tagged Lifeact (red). Simultaneous time-lapse imaging of AP2A1 and integrin β1 along the length of stress fibers was recorded with confocal microscopy with a 30-sec interval time for 10 min. To determine the speed of protein movement, we tracked AP2A1 and integrin β1 using TrackMate plugin in the Fiji/ImageJ software.

### 2.18 Statistical analysis

Each experiment was independently repeated at least three times, and results are expressed as means ± standard deviations. Statistical analyses were performed using GraphPad Prism (v.9.0.0) software. The statistical significance of differences among varying passage numbers and AP2A1 knockdown conditions (siControl, siAP2A1(1), siAP2A1(2)) was determined using a one-way analysis of variance (ANOVA), followed by Dennett’s multiple comparison test. The significance between two experimental groups was analyzed using unpaired two-tailed Student’s *t*-test. Statistical significance is described as follows: *, *p* ≤ 0.05; **, *p* ≤ 0.01; ***, *p* ≤ 0.001; and, ****, *p* ≤ 0.0001.

## 3. Results

### 3.1 Characterization of replicative senescent fibroblasts

Primary human foreskin fibroblast HFF-1 cells were expanded by serial passaging until passage 30 (P30) to induce replicative senescence. Cells at P10 were designated as young control, and those at P20 and P30 were categorized as the adult and aged cell groups, respectively. We found that cells at late passage exhibited an enlarged and flattened cell morphology (Fig. 1A). The area of aged cells (P30) increased by 6.6-fold compared to young control, whereas the nucleus size increased by 1.5-fold (Fig. 1A-B and Suppl. Fig. S1A). With increasing the passage number, cells also underwent reduction in the aspect ratio from a spindle shape to an irregularly round one (Fig. 1C). We examined some senescence markers to determine how the large cell size is associated with senescence. As expected, enlarged cells at P30 exhibited senescence-specific characteristics, including a significant increase in SA-β-gal activity (Fig. 1D), delay in cell proliferation (Fig. 1E), upregulation of p53 and p21 (Fig. 1F), increase in cellular ATP levels (Supple. Fig. S1B), and downregulation of protein synthesis (Supple. Fig. S1C) in accordance with previous studies on fibroblast replicative senescence (Gorgoulis et al., 2005; Hayflick & Moorhead, 1961; Tsai et al., 2021). To test whether the cell shape alteration modulates cell migration, we performed wound healing assay. The results demonstrated that the migration ability of senescent cells was significantly lower than young cells (Supple. Fig. S1D). These results suggest that senescence is indeed induced to the fibroblasts by repeated passaging, exhibiting common replicative senescence phenotypes.

**Figure 1.**
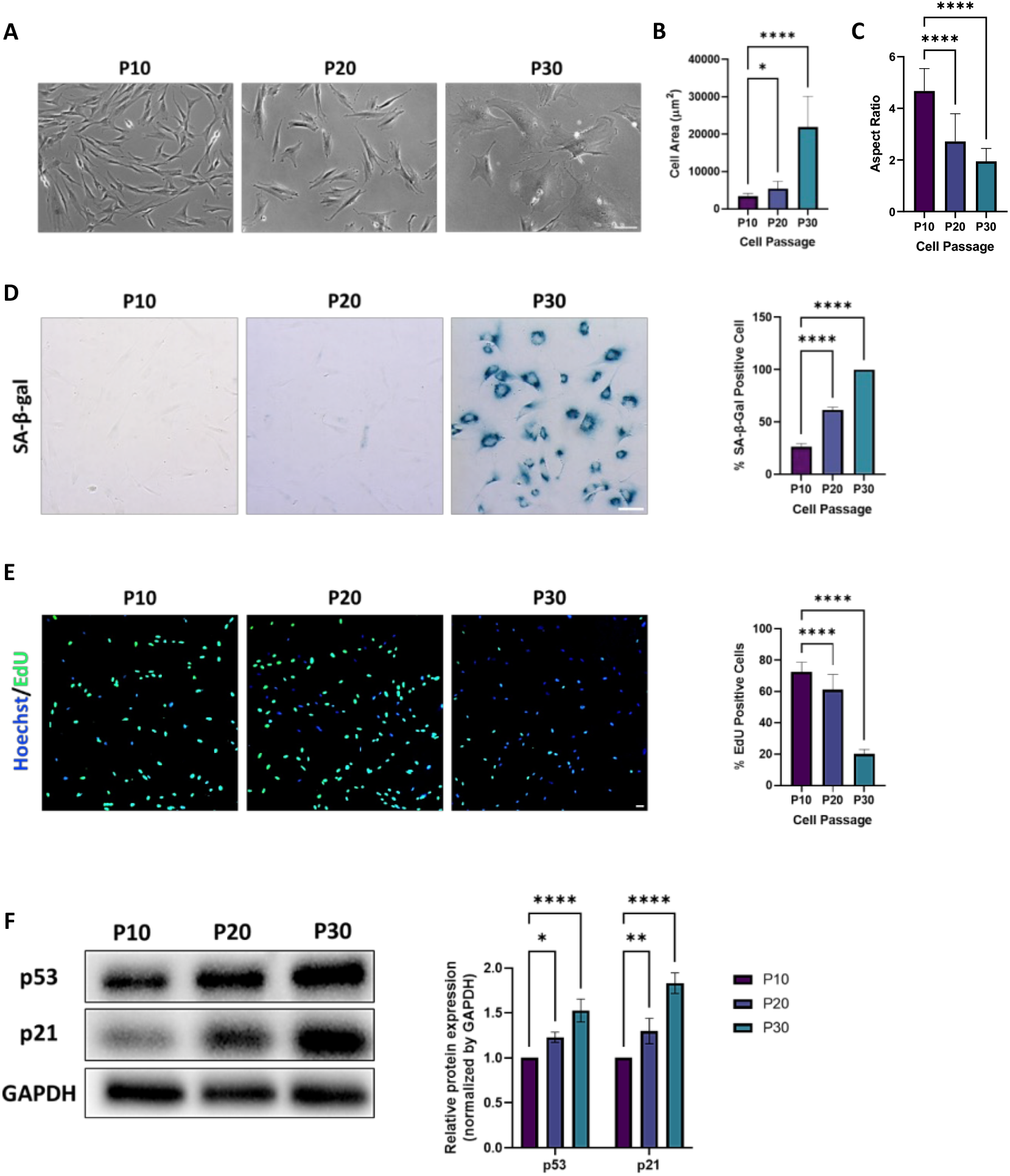
Characterization of replicative senescence of HFF-1 cells. (A) Phase-contrast microscopy of cells at P10, P20, and P30. Scale, 100 *µ*m. (B, C) Quantification of cell area (B; *n* = 30 cells, *N* = 3 independent experiments) and cell aspect ratio (C; *n* = 30 cells, *N* = 3 independent experiments). (D) SA-β-gal staining images and quantification (*n* = 10 random fields, *N* = 3 independent experiments). Scale, 50 *µ*m. (E) EdU staining images and quantification to evaluate cell proliferation (*n* = 10 random fields, *N* = 3 independent experiments). Scale, 50 *µ*m. (F) Representative western blot of senescence markers, p53 and p21, and quantification normalized by GAPDH (*N* = 3 independent experiments).

### 3.2 Replicative senescence alters stress fiber organization

We next investigated how the above change in cell morphology is associated with stress fiber organization. Phalloidin staining showed that senescent cells express a greater number of stress fibers per cell alongside a significant increase in the thickness compared to control (Fig. 2A and 2C-D). To evaluate the alignment of stress fibers within cells, the fluorescence images were analyzed by using 2D FFT (Supple. Fig. S2A). The overall distribution of fluorescence intensity, which represents the presence of stress fibers, is narrower in senescent cells compared to control, indicating that young cells form relatively randomly oriented stress fibers whereas senescent cells express more aligned forms. In addition, the expression level of typical stress fiber-associated proteins was determined by western blot analysis (Fig. 2B). We found that non-muscle myosin II (NMMIIa) and α-actinin are upregulated, while β-actin behaves as a house-keeping protein during the progression of senescence, thus consistent with our observations that the thickness of individual stress fibers is increased in senescent cells (Fig. 2E).

**Figure 2.**
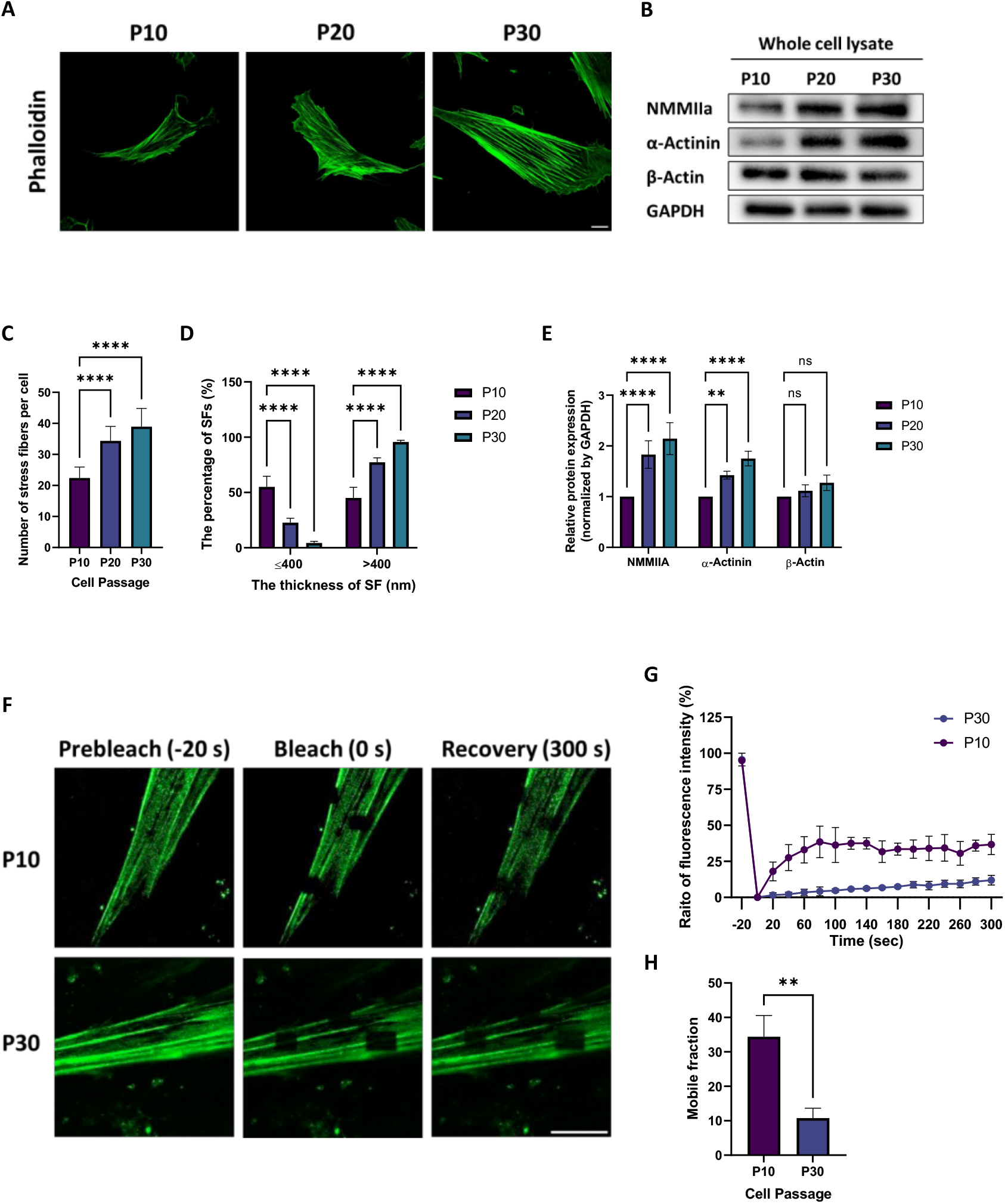
Response of stress fibers to replicative senescence. (A) Stress fibers in HFF-1 cells at P10, P20, and P30. Scale, 20 *µ*m. (B) Western blot of major stress fiber-associated proteins, NMMIIa, α-actinin, and β-actin (*N* = 3 independent experiments). (C) The number of individual stress fibers (SFs) per cell (*n* = 30 cells for each group). (D) The ratio of the number of individual stress fibers with a thickness of ≤ 400 nm or of > 400 nm to the whole population at P10, P20, and P30 (*n* ≥ 300 stress fibers for each group from *N* = 3 independent experiments). (E) Quantification of the western blot.

Next, we conducted fluorescence recovery after photobleaching (FRAP) to evaluate the turnover or the molecular stability of stress fibers in P10 and P30 cells, in which EGFP-MLC was monitored (Fig. 2F). The rate of MLC fluorescence recovery in senescent cells was extremely slower than in young cells (Fig. 2G). The recovery curves were quantified at the end timepoint (*t* = 300 s) to determine the mobile fraction, which was significantly lower in senescent cells (Fig. 2H). These results indicate that the structure of stress fibers is more stabilized in senescent cells.

### 3.3 Upregulation of AP2A1 upon fibroblast replicative senescence

Previous proteomic data revealed that AP2A protein expression is upregulated in the stress fiber fraction in response to fibroblast replicative senescence (Liu et al., 2022). To confirm this result with western blot, proteins from whole cell lysates and extracted stress fibers were analyzed at P10, P20, and P30. AP2A1 expression increased in both whole cell lysates and extracted stress fibers with the increase in passage number (Fig. 3A-D), which agrees with the proteomic analysis. Immunofluorescence staining was also performed to investigate the localization of endogenous AP2A1 in cells (Fig. 3E). Senescent cells displayed clearer fibrous patterns of AP2A1 compared to young cells that express more random patterns. 2D FFT image analysis was performed to quantify this feature, in which consistent geometric alignment of AP2A1 was indeed observed (Supple. Fig. S2B) in a manner similar to actin-labeled stress fibers (Supple. Fig. S1A). Quantitative colocalization of AP2A1 and stress fibers in confocal images was analyzed, showing a greater value of Pearson’s correlation coefficient in senescent cells (Fig. 3F). 3D-super-resolution SIM was conducted to observe detailes, and AP2A1 protein was found to localize just above or attach to the side of individual stress fibers (Fig. 3G). These observations suggest that AP2A1 works along the surface of stress fibers in senenscent cells.

**Figure 3.**
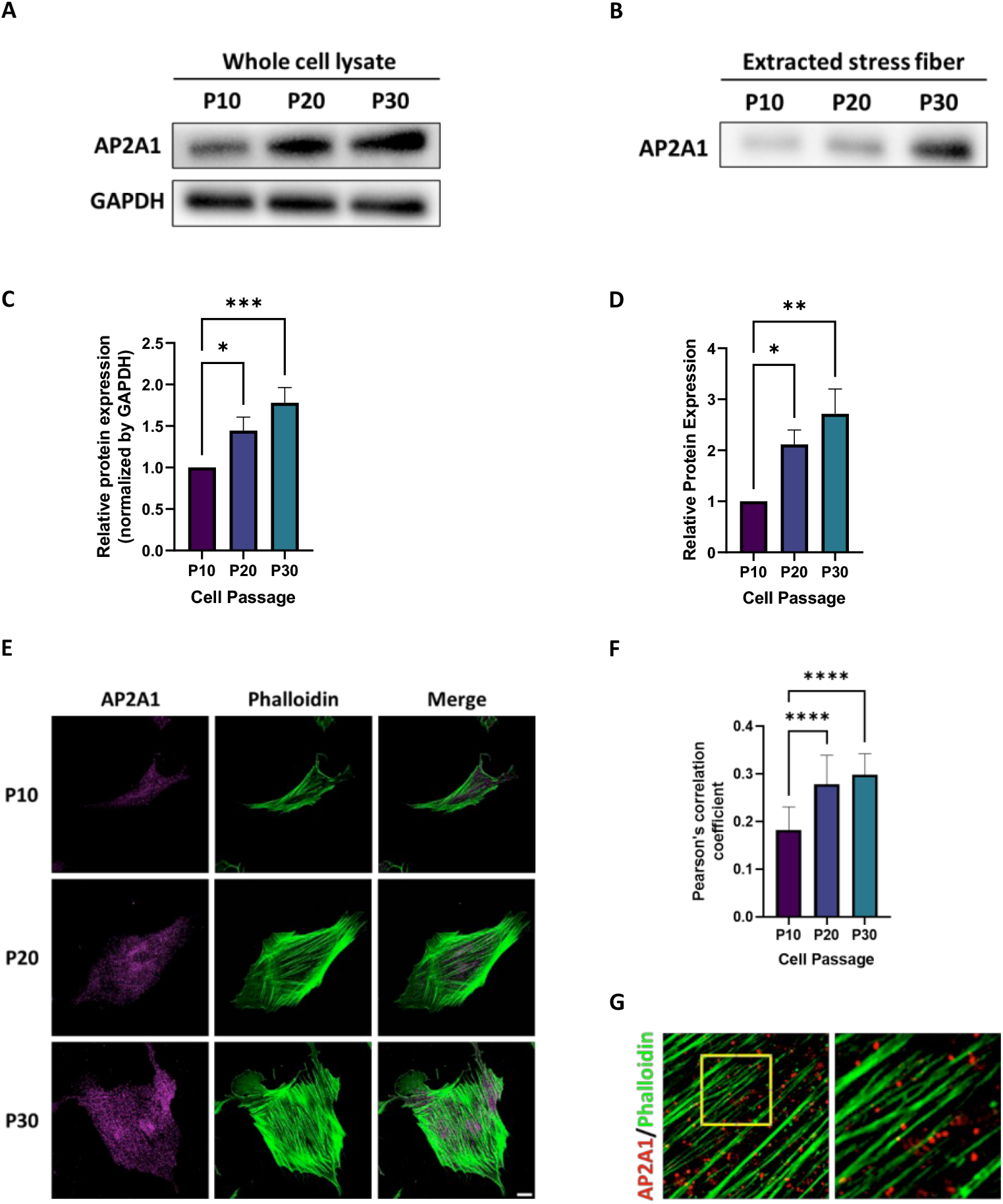
Increased AP2A1 protein expression in senescent cells. (A, B) Western blot of AP2A1 in the whole cell lysate (A; *N* = 3 independent experiments) and in extracted stress fibers (B; *N* = 3 independent experiments). (C, D) Quantification of the expression of AP2A1 in the whole cell lysate (C) and extracted stress fibers (D). (E) Immunostaining of endogenous AP2A1 (magenta) and actin filaments in stress fibers (phalloidin, green). (F) Pearson’s correlation coefficient to evaluate colocalization of AP2A1 and actin filaments (*n* = 20 cells). (G) 3D reconstructed SIM image of AP2A1 and actin filaments in stress fibers (phalloidin). The rectangles in yellow are magnified to show the detail.

As AP2A1 is known to be a major protein for vesicle formation in CME (Shin et al., 2021), we next investigated whether the level of AP2A1 affects endocytosis by using transferrin endocytosis assay. The intarnalization of fluorescencently labeled transferrin was imaged, and quantitatification showed that the total cellular uptake increased with senescence (Supple. Fig. S3A-B). To establish the role of AP2A1 in endocytic trafficking, we analyzed the colocalization of AP2A1 and transferrin. Quantification of the fluorescence images showed AP2A1 localizes with transferrin in both young and senescent cells with a high value of Pearson’s correlation coefficient of more than 0.7, and no significant difference was observed between the two cell groups (Supple. Fig. S3C).

### 3.4 AP2A1 regulates cellular senescence

The concomitant upregulation of AP2A1 and senescence markers upon increased passage number led us to investigate the effects of knockdown and overexpression of AP2A1 on the phenotypes associated with cellular senescnece. First, we used two different small interfering RNA (siRNA) sequences to deplete AP2A1 in senescent cells of P30. Upon silence of AP2A1, immunofluorescence showed that senescent cells exhibited, compared to control, a smaller cell area with a significant decrease in the thickness and the number of individual stress fibers per cell (Fig. 4A-C). Western blot showed that the expression of the major stress fiber-associated proteins, namely NMMIIa and α-actinin, was significantly reduced compared with control; meanwhile, β-actin remains unchanged (Fig. 4D-E). Interestingly, we found that the replicative senescence markers, namely SA-β-gal activity (Supple. Fig. S4A-B) and protein levels of p53 and p21, were all downregulated by the knockdown of AP2A1 (Fig. 4F-G), while cell proliferation was enhanced (Supple. Fig. S4C-D). AP2A1 silencing was also found to significantly reduce transferrin receptor uptake (Supple. Fig. S4E-F).

**Figure 4.**
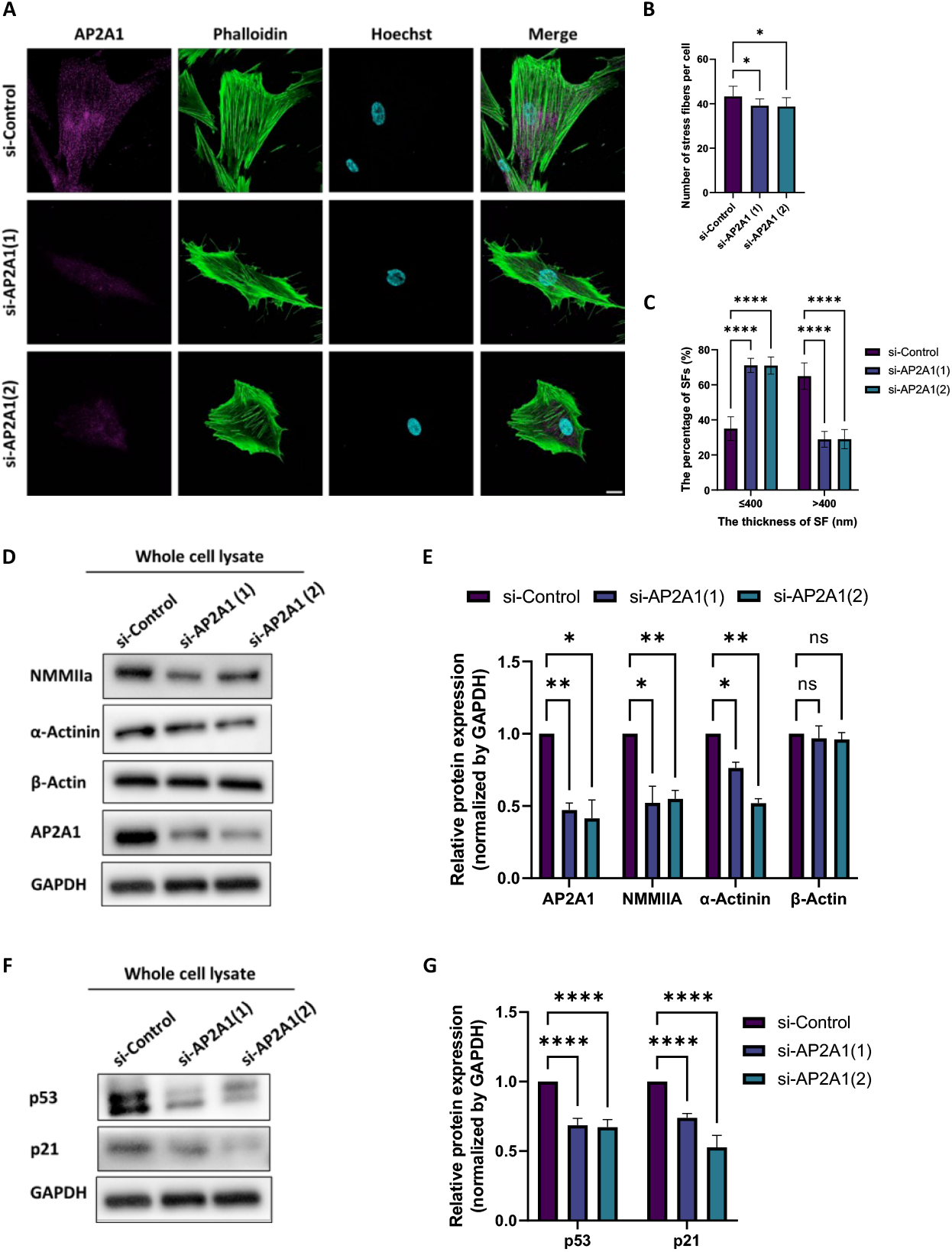
AP2A1 knockdown reduces stress fibers and senescence markers. (A) Immunostaining of AP2A1 (magenta), actin filaments in stress fibers (phalloidin, green), and nuclei (Hoechst33342, blue) in senescent P30 cells with si-control or si-AP2A1 transfection. Scale, 20 *µ*m. (B, C) Quantification shows the number of individual stress fibers (SFs) per cell (B) and the thickness (C) under the knockdown (*n* = 20 cells for each group from 3 independent experiments). (D, E) Western blot shows that AP2A1 knockdown in P30 cells led to a reduced expression of NMMIIa and α-actinin. (F, G) Western blot shows that the expression of p53 and p21 was both reduced by AP2A1 knockdown. The blots are representative of *N* =3 independent experiments.

In contrast, the overexpression of AP2A1 in young cells of P10 induced an increase in cell area as well as in the number and thickness of individual stress fibers (Fig. 5A-C) in a manner consistent with the upregulation of NMMIIa and α-actinin; meanwhile, β-actin remains unchanged in expression (Fig. 5D-F). The protein expression levels of p53 and p21 (Fig. 5E-G) were upregulated upon the AP2A1 overexpression. Given the opposite effects driven by AP2A1 knockdown and overexpression that suppress and promote the typical senescence phenotypes, respectively, these data suggest that the expression of AP2A1 regulates the extent of cellular senescence.

**Figure 5.**
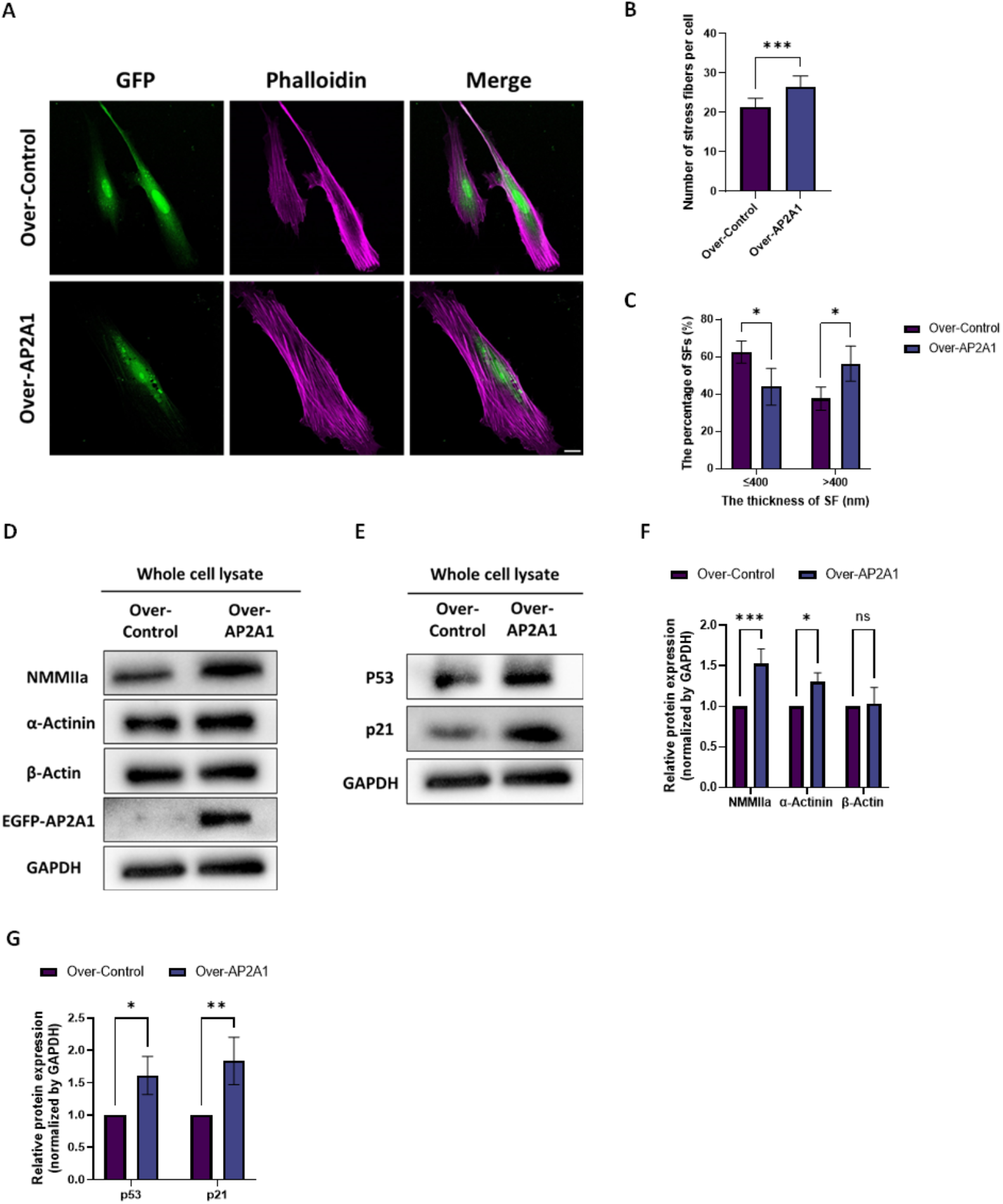
Overexpression of AP2A1 in young cells promotes senescence. (A) Fluorescence images of EGFP (AP2A1, green) and phalloidin (actin filaments, magenta) for cells with EGFP-AP2A1 overexpression. Scale, 20 *µ*m. (B, C) Quantification shows the number of individual stress fibers per cell (B) and the thickness (C) under the overexpression (*n* = 30 cells for each group). (D, E) Western blot of major stress fiber-associated proteins and AP2A1 (D) and senescence markers (E). An AP2A1 antibody was used for the detection of AP2A1. (F, G) Quantification of the western blot. The blots are representative of *N* = 3 independent experiments.

### 3.5 Senescence promotes focal adhesions

Given that cells spread on the extra cellular matrix (ECM) via integrin and form focal adhesions (FAs) to obtain a desirable state (Berrier & Yamada, 2007; Chao & Kunz, 2009), it is hypothesized that increased cell area in senescence is accompanied by enhanced structural maturity of FAs. To examine this relationship, we examined by immunostaining the expression of FA-associated proteins, vinculin and paxillin, in both young and senescent cell groups. Cells of each group were divided into 2 regions, edge and non-edge. Here, edge was defined as the intracellular regions within a distance of 5 *µ*m from the outline of the cells, and the rest of the intracellular regions was defined as non-edge. The number and size of individual FAs were quantified for each cell group. The quantification showed that in young cells both vinculin and paxillin are preferentially distributed around the cell edge, whereas in senescent cells those proteins were rather located at the cell center (Fig. 6A-D and Supple. Table S1). Stress fibers in young and senescent cells likewise tend to be located around the periphery and central regions of the cells, respectively (Supple. Fig. S5A-B). The size (Fig. 6C-D) and the number of FAs per cell (Supple. Table S1) were significantly higher in senescent cells. These results suggest that cellular senescnece exhibiting increased cell area is accompanied by the maturity of FAs.

**Figure 6.**
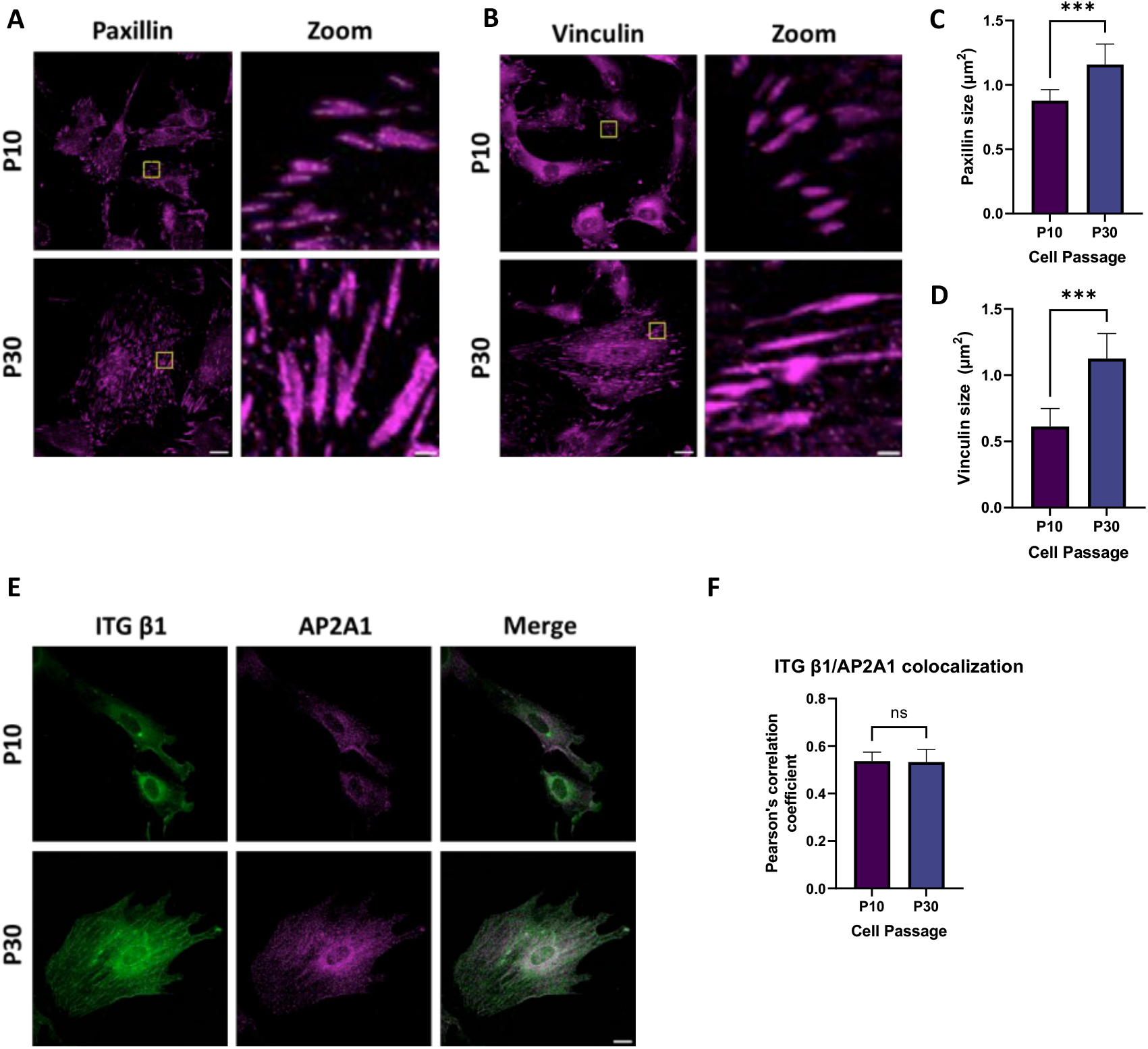
Focal adhesion maturation in response to senescence. (A, B) Immunostaining of paxillin (A) and vinculin (B) in P10 (control) and P30 (senescence) cells. Scale, 20 *µ*m. (C, D) Quantification of the area of individual FAs determined by the paxillin (C) and vinculin (D) images (*n* = 30 cells for each group from *N* = 3 independent experiments). (E) Representative single channel and merged images of integrin β1 and AP2A1 in P10 (young) and P30 (senescence) cells. Scale, 20 *µ*m. (F) Quantification of the colocalization of integrin β1 and AP2A1 (*n* = 20 cells for each group *N* = 3 independent experiments).

We further investigated how the cell adhesions to the ECM is strengthend during senescence by using trypsin-based cell detachment assay. The time required for de-adhesion with trypsinization was significantly delayed in senescent cells compared to young cells (Supple. Fig. S6A-B). Thus, FAs in senescent cells are not only augmented in size and molecular expression but also strengthened in terms of the anchorage to the ECM.

### 3.6 AP2A1 is required for integrin β1 translocation

Endocytosis of vesicles carrying integrin β1 via CME has been proposed to be a mechanism to supply integrin for newly forming FAs (Gu et al., 2011; Pankov et al., 2000). Based on the above results that FAs are strengthened upon senescence, we hypothesized that AP2A1 is involved in translocation of integrin β1 along the length of stress fibers. To test this, we first performed double immunofluorescence staining of AP2A1 and integrin β1. Partial localization of integrin β1 and AP2A1 was observed with a Pearson’s correlation coefficient of over 0.5, while there is no significant difference in the extent between young and senescent cell groups (Fig. 6F-G). Integrin β1 may be recycled back to the plasma membrane via endosomes or targeted for degradation at lysosomes, namely autophagy (Kenific et al., 2016). In this regard, we analyzed the colocalization of integrin β1 with the lysosome marker, LAMP1. We observed a decrease in integrin β1 overlap with LAMP1 upon senescence (Supple. Fig. S7A). These results inspired us to consider that senescent cells tend to promote the recycle of internalized integrin β1 back to the plasma membrane as a potential new resource to maintain FAs, while young cells rather allow internalized integrin β1 to be transported to degradative lysosomes.

To analyze the involvement of AP2A1 in integrin β1 transportation, living cells transfected with EGFP-AP2A1 and mRuby-Lifeact were imaged by time-lapse confocal microscopy at an interval of 30 sec for 10 min. The fluorescence of individual AP2A1 aggregations was tracked to demonstrate that AP2A1 moves along the length of stress fibers (Movies 1-2; not included for bioRxiv version). The speed of the AP2A1 movement was significantly faster in young cells compared with senescent cells (Fig. 7A-B). Using the same approach, we also observed integrin β1 translocation along stress fibers in cells transfected with mVenus-integrin β1 and mRuby-Lifeact (Movie 3-4; not included for bioRxiv version). The integrin β1 movement was substantially faster in young cells, consistent with the observation of the AP2A1 dynamics (Fig. 7C-D). These data support our hypothesis that integrin β1 contained in a vesicle with AP2A1 is translocated by CME along stress fibers to FAs where integrin β1 is clustered (Fig. 7E), and the turnover rate of integrin β1 is slowed in senescent cells as the associated stress fibers are stabilized (Fig. 2F-H).

**Figure 7.**
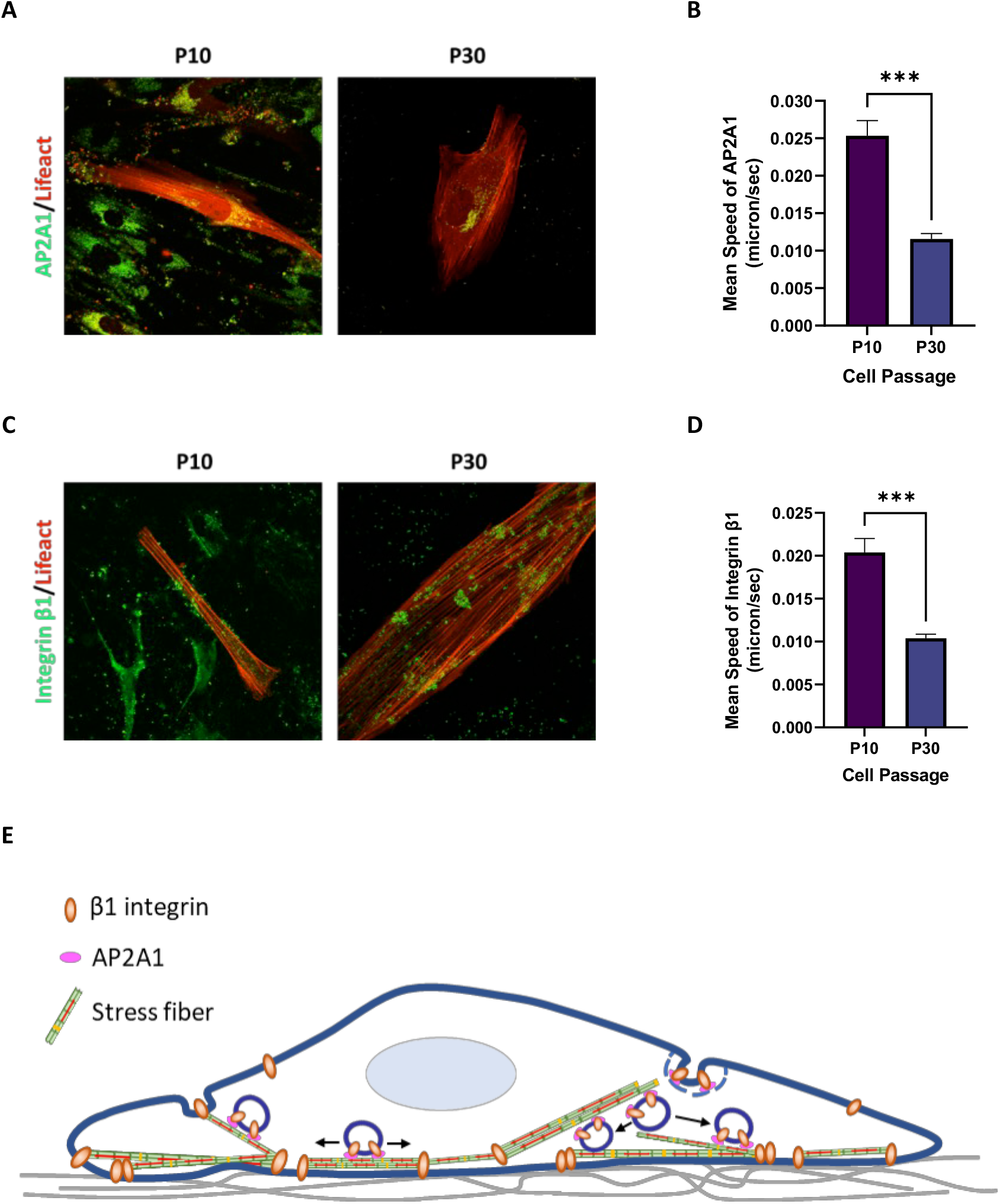
Analysis of protein movement along stress fibers. (A) Snapshot of Video 1, in which EGFP-AP2A1 (green) and mRuby2-Lifeact (red) were expressed in P10 (young) and P30 (senescence) cells to observe the movement. (B) Quantification of the speed of AP2A1 movement (*n* = 30) (C) Snapshot of Video 2, in which mVenus-integrin β1 (green) and mRuby2-Lifeact (red) were expressed in P10 (young) and P30 (senescence) cells to observe the movement. (D) Quantification of the speed of integrin β1 movement (*n* = 30). (E) Schematic of integrin β1 translocation along stress fibers via CME.

## 4. Discussion

The function of organisms progressively declines during the lifespan due to overaccumulation of senescent cells in multiple tissues, leading to age-related diseases (López-Otin et al., 2013; van Deursen, 2014). Many researches have attempted to develop therapeutic drugs targeting senescent cells, and cultured human fibroblasts are often used in those studies as an in vitro model to induce cellular senescence (Hayflick, 1965; Hayflick & Moorhead, 1961; Tigges et al., 2014). In the present study, we established a replicative senescence model of HFF-1 cells by serial passage. Cells at passage 30 exhibited typical senescence characteristics, e.g., an enlarged and flattened cell shape (Fig. 1A), increased SA-β-galactosidase activity (Fig. 1D), and upregulated p53 and p21 (Fig. 1F), thus consistent with previous studies (Collado & Serrano, 2010; Cristofalo & Pignolo, 1996; Dimri et al., 1995). Many studies on senescence have also reported that cell size increases gradually during replicative senescence (Angello et al., 1987; Kim et al., 2013), and alteration in cell size contributes to decline in cell function, proliferation, and protein synthesis (Lloyd, 2013; Neurohr et al., 2019). However, there is little understanding of the mechanism behind the significant changes in stress fibers as well as focal adhesions that are known to be elicited upon replicative senescence.

We found that major senescence phenotypes are indeed induced to the human fibroblasts with repeated passage (Fig. 1), in which stress fibers were increased in the total number and thickness (Fig. 2). In the aged stress fibers, higher expression levels of actin-binding proteins, specifically, non-muscle myosin II and α-actinin, were associated compared to control young stress fibers. The turnover rate of stress fibers, assessed by the FRAP experiments, was higher in senescent cells than in young cells, suggesting that stress fibers are stabilized upon senescence. Given the association of stress fibers with cell migration (Lehtimäki et al., 2021), we investigated the effect of expressing the stabilized stress fiber organization on cell migration and found that it slows cell migration (Supple. Fig. S1D).

Toward better understanding of senescence-related change in stress fiber organization, our group previously conducted proteomic analysis on HFF-1 cells with or without replicative senescence, identifying that 63 different proteins are upregulated along stress fibers in response to the induced senescence (Liu et al., 2022). Approximately two-thirds of them are not known for the function related to stress fibers as well as how it plays a role in senescence, and therefore among them here we focused on AP2A1. We confirmed the upregulation of endogenous AP2A1 along stress fibers in the same human fibroblasts HFF1 cells upon senescence progression by western blotting and immunostaining (Fig. 3A-F). AP2A1 has been implicated as a potential biomarker among patients with ovarian cancer (Jiao et al., 2021), pediatric medulloblastoma (Huang et al., 2020), and Alzheimer’s disease (Raj et al., 2018; Tian et al., 2013). Although AP2A1 was thus suggested to be a pathological marker for aging-associated diseases, the underlying mechanisms were yet to be identified (Shin et al., 2021). In this regard, we demonstrated that AP2A1 plays a role in cellular senescence by modulating the size of individual stress fibers and FAs. The aberrant expression of AP2A1 in senescent cells may results from a higher total cellular uptake surface area (Wang et al., 2016) and an increase in travel time (Miettinen et al., 2017) and distance for endocytosed vesicle transportation, which is further discussed below.

We demonstrated that AP2A1 knockdown in senescent cells decreases not only senescence markers but also cell size and stress fiber abundance (Fig. 4). In addition, the overexpression of AP2A1 in young cells promotes cell senescence with an increased cell size and stress fiber abundance (Fig. 5). Thus, AP2A1 is involved in morphological alteration and stress fiber reorganization in senescent cells. It is therefore interesting to identify the molecule that is transported by AP2A1 to control senescence phenotype.

Cell adhesion, spreading, and migration on the substrate are modulated by the integrin-mediated formation of FA complexes (Hakkinen et al., 2011; Kim & Wirtz, 2013; Yeung et al., 2005). Our observation that extensively spread senescent cells express highly mature FAs with a large size (Fig. 6A-D) led us to focus on integrin endocytic trafficking. The integrin αβ heterodimers play a crucial role in cell-matrix attachment, and the majority of integrin receptors contain the β1 subunit (Moreno-Layseca et al., 2019). Under certain conditions, integrin adhesions are highly dynamic, undergoing continuous cycles of assembly and disassembly to facilitate the regulation of cell adhesion. The main integrin internalization route is via CME (Bridgewater et al., 2012; Paul et al., 2015). Given these reports, our observations on the colocalization of integrin β1 and AP2A1 required for the CME suggest that AP2A1, becoming associated with stress fibers in senescent cells, is involved in vesicle-bound integrin β1 transportation in the senescent state. In fact, it has been reported that in fibroblasts fibronectin-bound integrin α5β1 translocates along stress fibers (Kechagia et al., 2019). The movement of vesicles over a long range along stress fibers is mediated by myosin 5b (Schuh, 2011). Other studies demonstrated translocation of vesicles containing integrin from the intracellular space to the extending tip of cell adhesions by stress fiber-dependent translocation (Kawakami et al., 2001; Lawson & Maxfield, 1995). All heterodimers containing integrin β1 have thus been shown to be potentially endocytosed via the same mechanism (Arjonen et al., 2012; Calderwood et al., 2003). In addition, autophagy, namely a lysosome-dependent cellular degradation process, works as another regulator of integrin-mediated cell adhesion (Vlahakis & Debnath, 2017). Particularly, suppression of autophagy has been implicated in cellular senescence (Kim et al., 2013). Thus, integrin can be recycled back to the plasma membrane after internalization or degraded in lysosomes via autophagy (Kenific et al., 2016). We observed downregulation of integrin β1 degradation (Fig. 6E-F and Supple. Fig. S7A-B) and stabilized translocation of integrin along stress fibers (Fig. 7C-D) in senescent cells, supporting enhanced integrin recycling rather than degradation. The integrin translocation along stress fibers may be efficient to maintain FAs in senescent cells, instead of transporting the molecules in a random diffusion-limited manner, given the greatly enlarged cell volume and resulting increase in travel time and distance for endocytosed vesicle transportation (Fig. 7E).

Taken together, we uncovered molecular mechanisms behind the cell enlargement in replicative senescence; specifically, aged fibroblasts exhibit enhanced integrin β1 translocation along stress fibers towards FAs, which requires AP2A1 to form the necessary vesicles. We propose a mechanistic account that individual FAs are significantly enlarged in senescent cells to assure firm anchorage to the substrate enough to sustain the huge cell architecture. Moreover, given the remarkable fibroblast responses to the expression level, the present study suggests that AP2A1 may serve as a new senescence marker as well as a therapeutic target for age-related diseases.

## Acknowledgments

The authors thank Natsuki Saito and Shinji Sakai (Osaka University) for their assistance in cellular ATP measurement. This study was partly supported by JSPS KAKENHI grant (21H03796) and JICA (Japan International Cooperation Agency).

## Supplementary information

**Figure S1.**
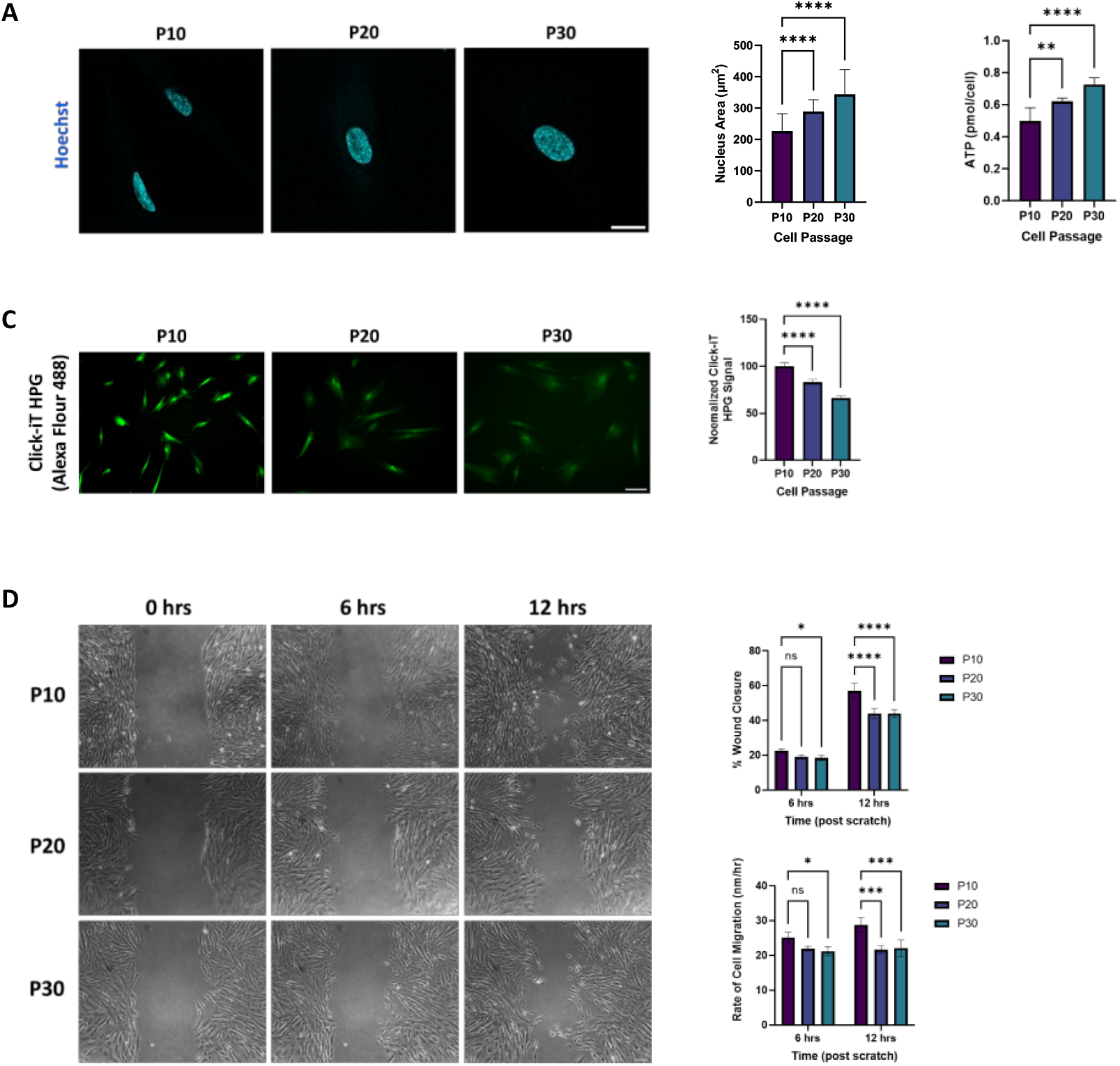
Confirmation of replicative senescent HFF-1 cells. (A) Cell nuclei at different passages were visualized using Hoechst 33342. Scale, 20 *µ*m. Quantification shows that average nucleus area is increased with passage number (*n* = 30 cells, *N* = 3 independent experiments). (B) Intracellular ATP levels at different senescence levels (*N* = 3 independent experiments). (C) Protein synthesis ability and its quantification (*n* = 10 random fields, *N* =3 independent experiments). (D) Cell migration (scratch wound healing assay). Representative images of the wound healing assay to evaluate migration rate at 6 and 12 h after scratch. Scale, 100 *µ*m. The right panels showed wound closure that was evaluated over the time by measuring the remaining area uncovered by the cells and expressed as a percentage of the cell-free area at 0 h and the wound healing rates (*N* = 3 independent experiments).

**Figure S2.**
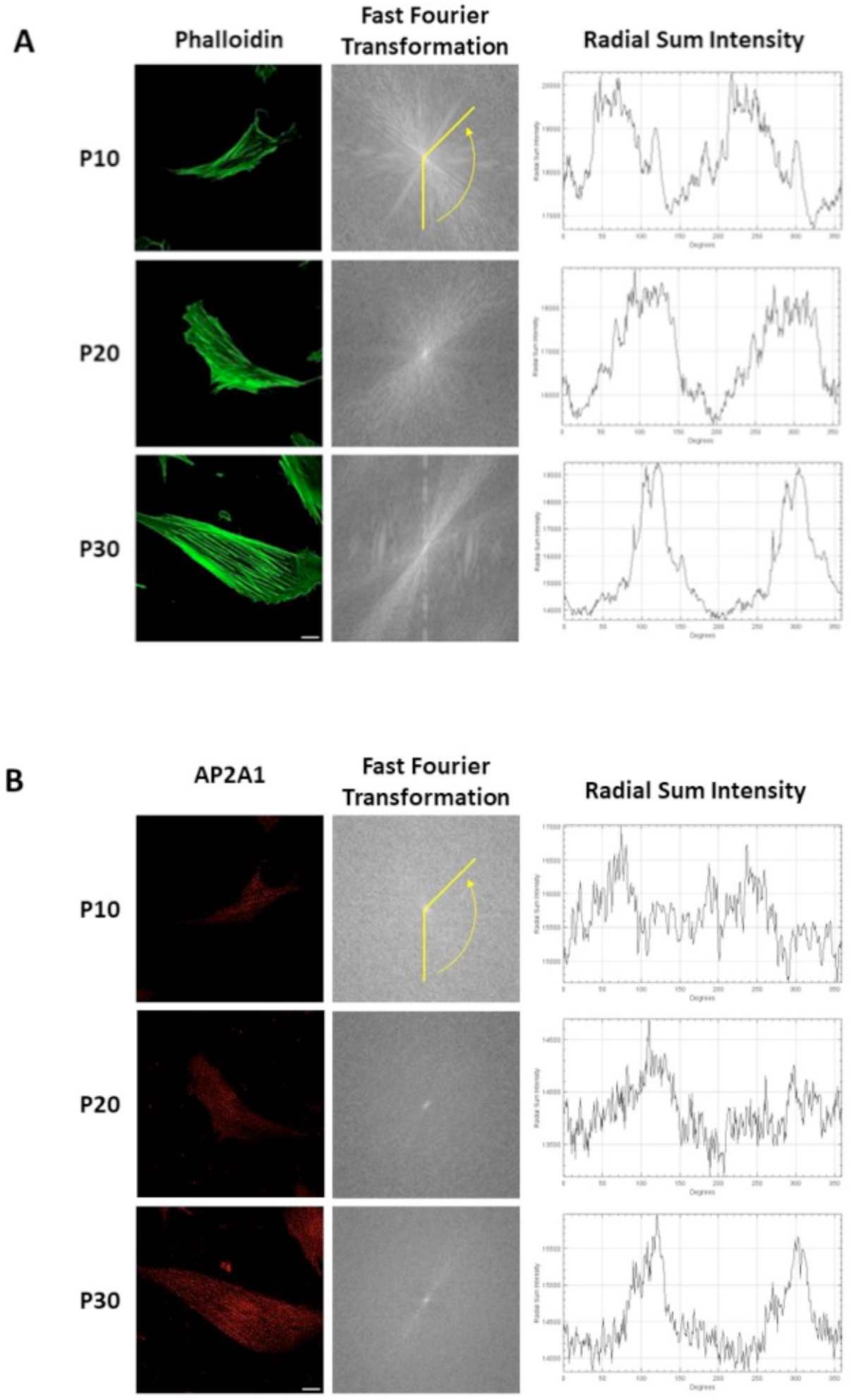
Analysis of structure alignment using 2D fast Fourier transform. (A) Analysis of stress fiber alignment for fluorescence images stained with phalloidin (green). Scale, 20 *µ*m. (B) Analysis of AP2A1 alignment for fluorescence images stained with anti-AP2A1 (red) in relation to stress fibers in (A).

**Figure S3.**
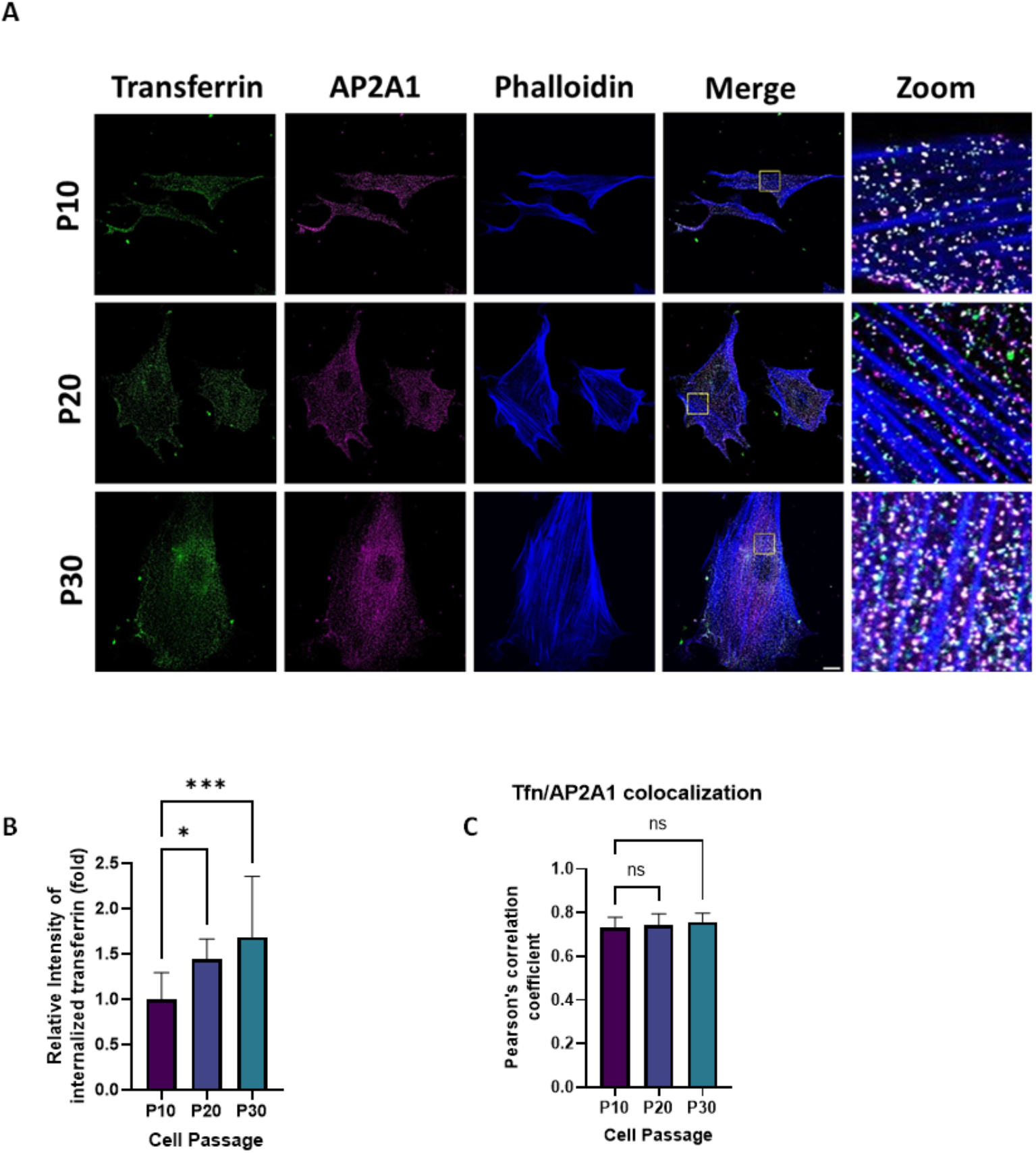
Senescence promotes transferrin uptake. (A) Transferrin endocytosis in HFF-1 cells at different levels of senescence. Transferrin uptake assay was performed using Alexa-Flour-488-conjugated transferrin (green). Cells were stained with anti-AP2A1 (magenta) and phalloidin (blue). Scale, 20 *µ*m. The areas with the yellow rectangle are magnified on the right. (B) Quantification of endocytosed transferrin in P20 and P30 compared to P10 (control). *n* = 10 cells for each group from *N* = 3 independent experiments. (C) Quantification of transferrin colocalization with the fibrous AP2A1 patterns (*n* = 20 cells).

**Figure S4.**
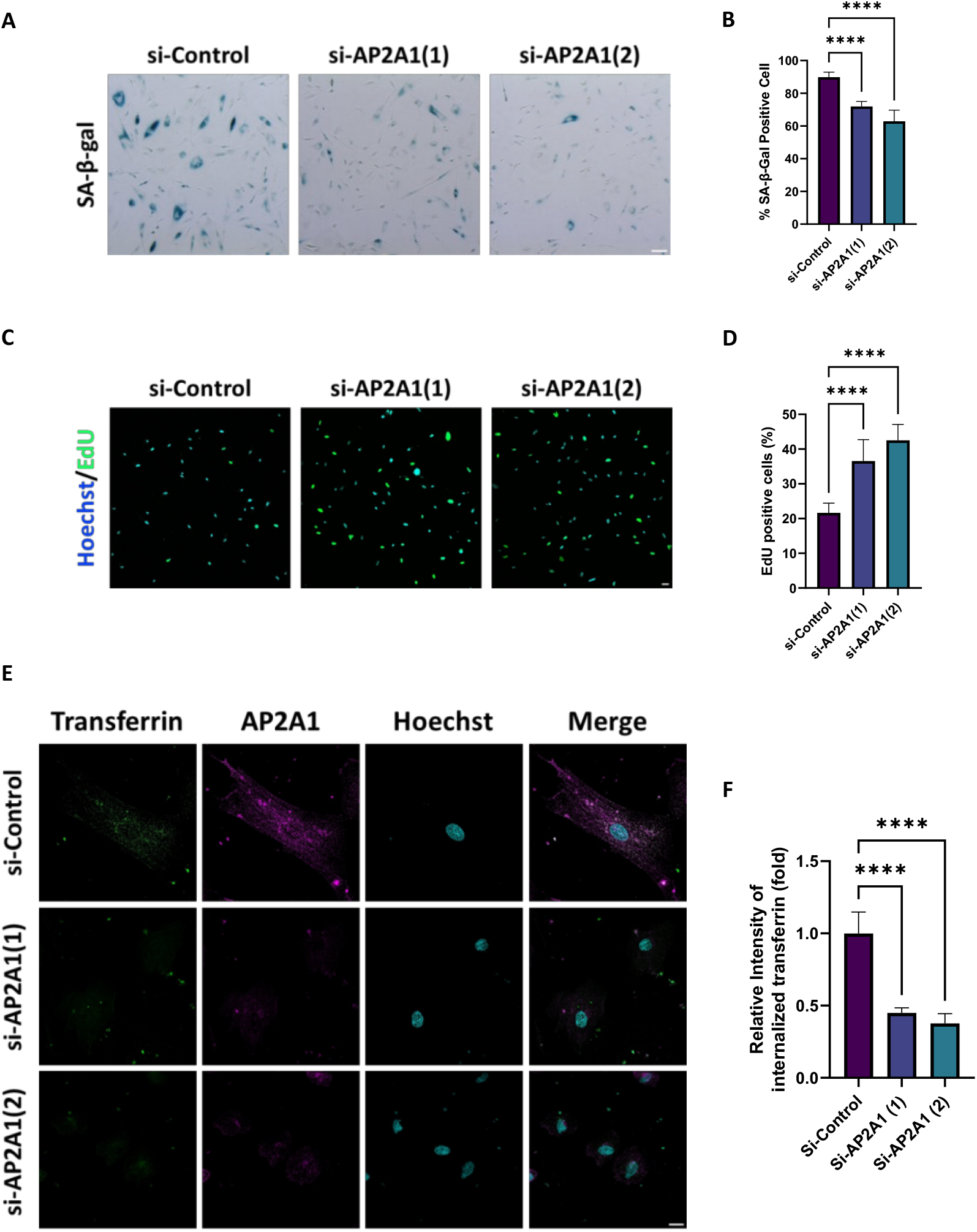
AP2A1 knockdown inhibits senescence phenotype and endocytosis ability. (A, B) AP2A1 knockdown in senescent cells reduced the level of SA-β-gal staining. Scale, 100 *µ*m. (*n* = 10 random fields for each group, *N* = 3 independent experiments). (C, D) Comparison of cell proliferation ability between senescent cells expressing siRNA for control or AP2A1 by EdU proliferation assay and its quantification. Scale, 20 *µ*m. (*n* = 10 random fields for each group, *N* =3 independent experiments). (E) Representative images of transferrin endocytosis in siRNA-treated cells (P30). Scale, 20 *µ*m. (F) Quantification of endocytosed transferrin in si-AP2A1-treated cells compared to Si-control-treated cells.

**Figure S5.**
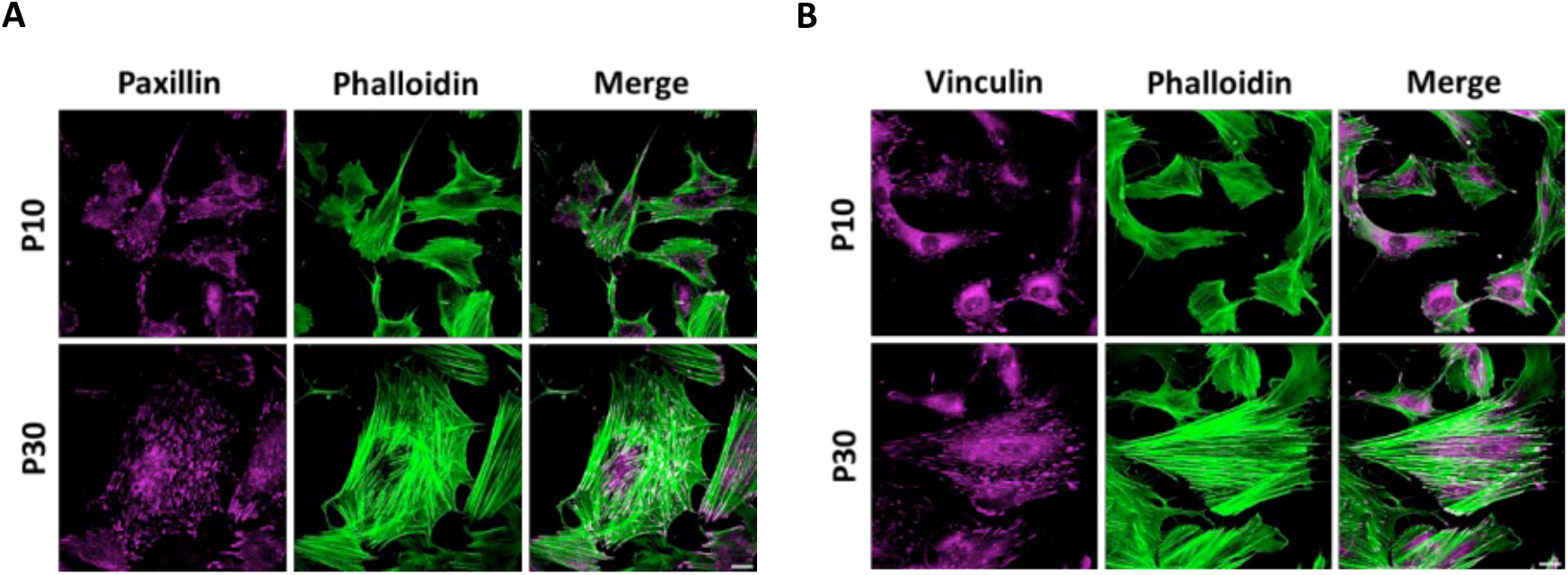
Immunofluorescence staining of focal adhesion-associated proteins. (A) paxillin (magenta) and (B) vinculin (magenta) and actin filaments in stress fibers (phalloidin, green). Scale, 20 *µ*m.

**Figure S6.**
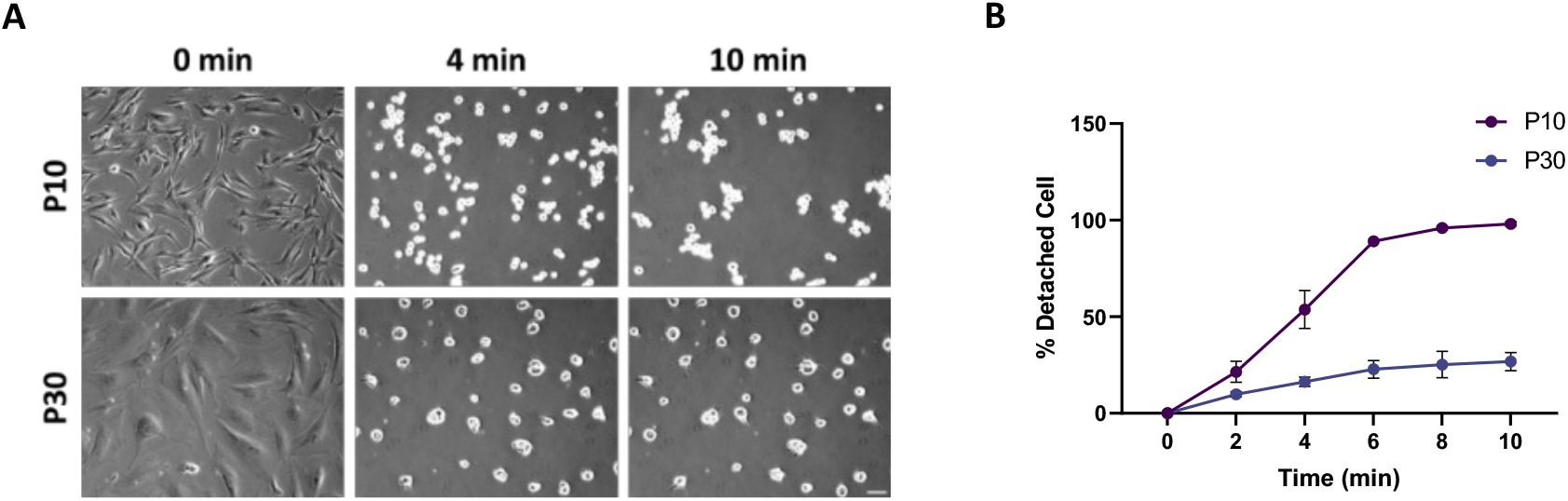
Influence of senescence on cell adhesion. (A) The strength of the cell attachment to the substrate in P10 (young) and P30 (senescence) cells was analyzed by trypsin-sensitive cell detachment assay. Scale, 100 *µ*m. (B) Quantitative results of the assay are shown as the percentage of detached cells (*N* = 3 independent experiments).

**Figure S7.**
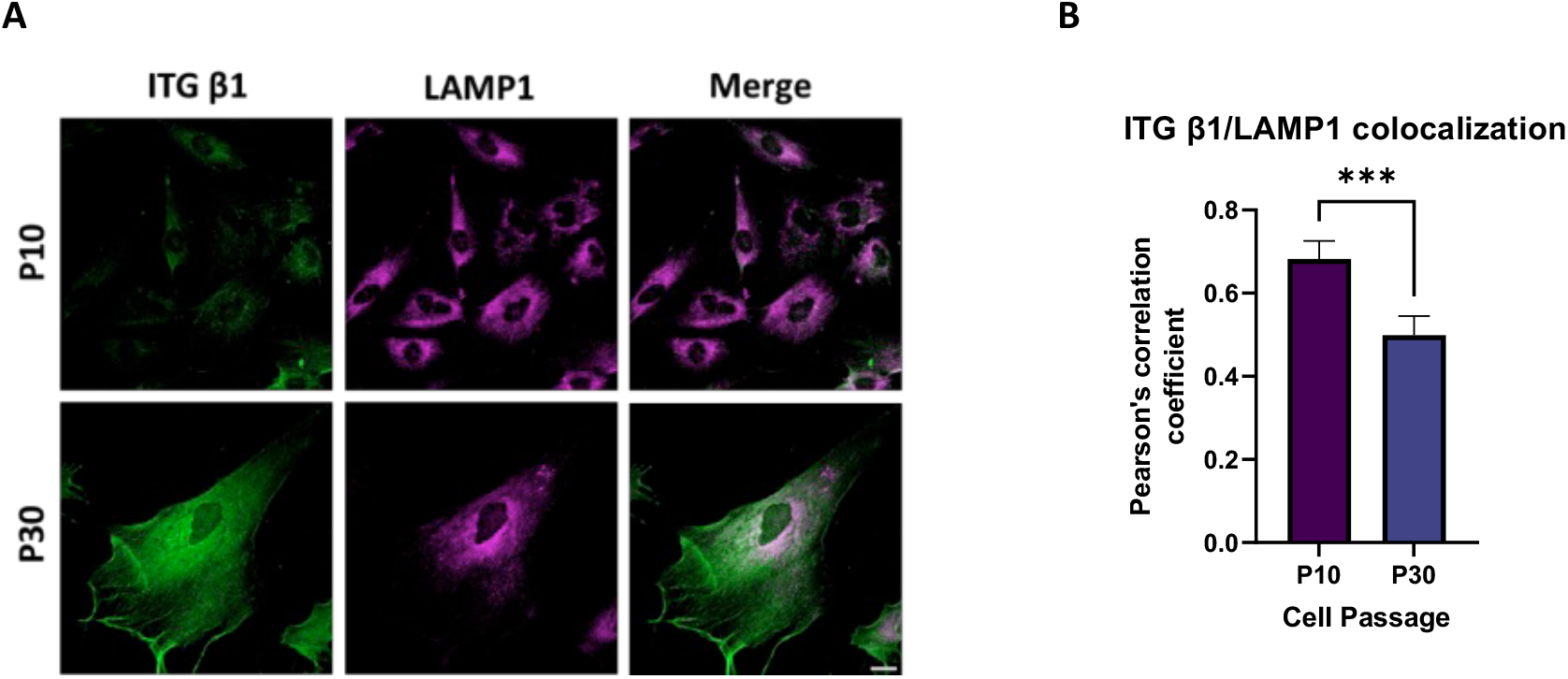
LAMP1 interacts with integrin β1 for the autophagy of integrin β1. (A) Representative single channel and merged images of integrin β1 (green) and LAMP1 (magenta) in P10 (young) and P30 (senescence) cells. Scale, 20 *µ*m. (B) Quantification of integrin β1 and LAMP1 colocalization (*n* = 20 cells).

**Table S1.** Focal protein distribution in P10 (young) and P30 (senescent) cells.

| **Cells** | **Paxillin** |  |  | **Vinculin** |  |  |
| --- | --- | --- | --- | --- | --- | --- |
|  | Total<br>number<br>per cells | Edge (%) | Non-edge (%) | Total<br>number per<br>cells | Edge (%) | Non-edge<br>(%) |
| P10 | 97.7±23.9 | 63.1 | 36.9 | 100±23.9 | 66.7 | 33.3 |
| P30 | 300±73.3 | 26.4 | 73.6 | 303.8±59.9 | 29.9 | 70.1 |

